# Polarized Mechanosensitive Signaling Domains Protect Arterial Endothelial Cells Against Inflammation

**DOI:** 10.1101/2023.05.26.542500

**Authors:** Soon-Gook Hong, Julianne W. Ashby, John P. Kennelly, Meigan Wu, Eesha Chattopadhyay, Rob Foreman, Peter Tontonoz, Patric Turowski, Marcus Gallagher-Jones, Julia J. Mack

## Abstract

Endothelial cells (ECs) in the descending aorta are exposed to high laminar shear stress, which supports an anti-inflammatory phenotype that protects them from atherosclerosis. High laminar shear stress also supports flow-aligned cell elongation and front-rear polarity, but whether this is required for athero-protective signaling is unclear. Here, we show that Caveolin-1-rich microdomains become polarized at the downstream end of ECs exposed to continuous high laminar flow. These microdomains are characterized by higher membrane rigidity, filamentous actin (F-actin) and lipid accumulation. Transient receptor potential vanilloid-type 4 (Trpv4) ion channels, while ubiquitously expressed, mediate localized Ca^2+^ entry at these microdomains where they physically interact with clustered Caveolin-1. The resultant focal bursts in Ca^2+^ activate the anti-inflammatory factor endothelial nitric oxide synthase (eNOS) within the confines of these domains. Importantly, we find that signaling at these domains requires both cell body elongation and sustained flow. Finally, Trpv4 signaling at these domains is necessary and sufficient to suppress inflammatory gene expression. Our work reveals a novel polarized mechanosensitive signaling hub that induces an anti-inflammatory response in arterial ECs exposed to high laminar shear stress.

## Introduction

Blood flow patterns in the aorta are defined by vessel geometry: the lower curvature of the aortic arch results in low/oscillatory flow, whereas the descending aorta experiences high laminar flow^1–4^. The luminal layer of endothelial cells (ECs) reveals a striking difference in collective cell morphology in these two regions where ECs lining the descending aorta are highly elongated^5, 6^. Endothelial cell alignment with flow induces a well-documented front-rear polarity with respect to the flow direction^7–11^, including preferential polarization of plasma membrane proteins NOTCH1 and VE-PTP to the downstream end of ECs^12–14^. ECs in the descending aorta also display a well-known anti-inflammatory and athero-protective phenotype^15, 16^, but the underlying mechanisms and connection to front-rear polarity are unknown. Furthermore, the role of plasma membrane polarization for anti-inflammatory signaling has not been described.

Nitric oxide (NO) production by endothelial nitric oxide synthase (eNOS) is a key anti-inflammatory response of arterial ECs to laminar flow^17–19^. Caveolin-1 microdomains play important roles in recruiting and activating eNOS^20, 21^. Furthermore, the ion channel Trpv4 associates with endothelial caveolae and contributes to Ca^2+^ dependent eNOS activation in pulmonary arteries^22–24^. Caveolae also play a role in the response of ECs to altered shear stress conditions^25, 26^, and chronic shear stress induces caveolae formation at the luminal surface of arterial ECs^27, 28^. Although caveolae have been observed to polarize in migrating cells^29, 30^, whether laminar flow affects the functional organization of caveolae is not known.

Caveolin-1, eNOS, Trpv4 and Ca^2+^ have each been reported for the maintenance of arterial vasodilation^31^, but this signaling pathway, its activation, and its physiological role in vascular inflammation are unclear. Here, we investigated the role of sustained flow on arterial EC signaling and found a distinct polarization of signaling activity, asymmetrically concentrated at the downstream end of cells. Polarization included membrane lipids, F-actin, and Caveolin-1 associated with Trpv4 channels. In the presence of high shear stress, localized Ca^2+^ oscillations via Trpv4 channels are sustained at these domains and contribute to anti-inflammatory gene expression via eNOS. Thus, our studies revealed a novel, focally restricted signaling domain in arterial ECs that is mechanosensitive and anti-inflammatory.

## Results

### Front-rear polarization of arterial endothelial cells

We considered that high laminar flow, which results in a greater than two-fold increase in EC aspect ratio in the descending aorta compared to the lower arch (Fig. S1a), affects caveolae organization. To test this hypothesis, we stained for components of caveolae, the plasma membrane-associated proteins Caveolin-1 and Cavin-1, in ECs lining the mouse descending aorta (Fig. 1a; Fig. S1c). We quantified protein distribution by segmenting each cell into three equal length regions (upstream, mid-body, and downstream) and found that the downstream end of descending aortic ECs displayed the highest levels of Cavin-1 and Caveolin-1.

**Fig. 1:**
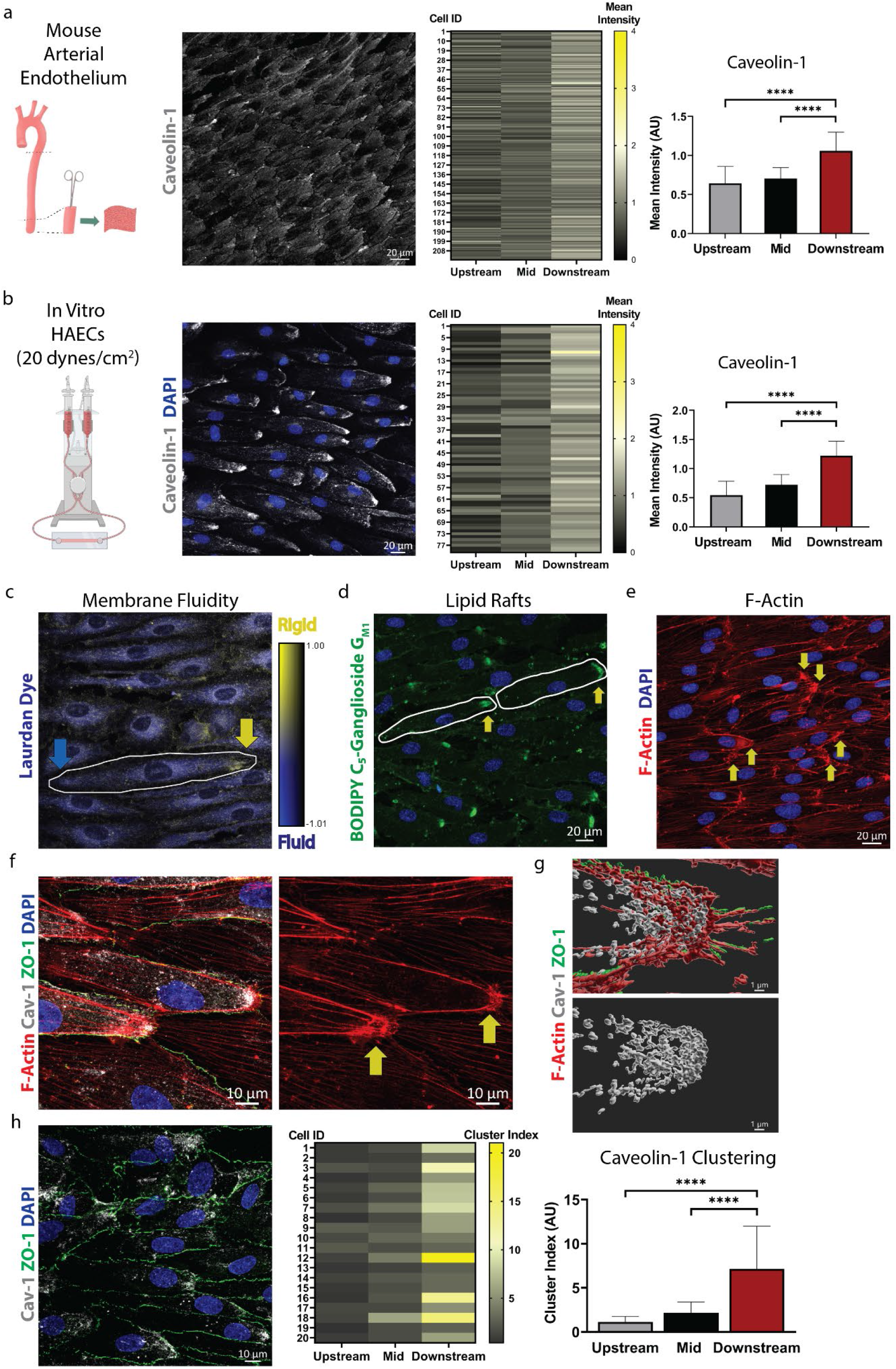
Membrane polarization and Caveolin-1-enriched microdomains in aortic ECs exposed to laminar flow. **a**, Confocal imaging of Caveolin-1 in the endothelium of wildtype mouse descending aorta. Individual cells were segmented into 3 equal-length regions (upstream, mid and downstream) and the staining intensity determined in each segment for 216 cells from n=3 aortas. Bar graph displays the mean ± SD with data analyzed by one-way ANOVA and post hoc Tukey’s multiple comparisons test; \*\*\*\**P* < 0.0001 (Upstream vs. Downstream; Mid vs. Downstream). For (b-h), confluent monolayers of HAECs were exposed to laminar shear stress (20 dynes/cm^2^) for 48 h then stained and imaged. **b,** HAECs were stained for Caveolin-1 and DAPI then segmented into 3 equal-length regions (upstream, mid and downstream) and Caveolin-1 staining intensity quantified for each subcellular region for 79 cells. Bar graph displays the mean ± SD with data analyzed by one-way ANOVA and post hoc Tukey’s multiple comparisons test. \*\*\*\**P* < 0.0001 (Upstream vs. Downstream; Mid vs. Downstream). **c**, Representative generalized polarization (GP) color-coded images from 2-photon imaging of cells stained with Laurdan dye to determine membrane fluidity, where higher GP indicates a more rigid membrane. Higher GP was observed at the downstream end (yellow arrow) compared to the upstream end (blue arrow). **d**, Staining live cells with BODIPY FL C_5_-Ganglioside G_M1_ showed polarized accumulation of fluorescent signal at the downstream end of cells (arrows). Nuclei are shown in blue. **e**, Fixed cells, stained for F-Actin, showed a higher density of staining at the downstream end of cells (arrows). Nuclei are shown in blue. **f**, Representative image of merged F-actin, Cav-1, junctional protein ZO-1 and nuclei at higher magnification highlights the presence of Cav-1 and F-Actin at the downstream end, and the formation of dense F-Actin web-like features (arrows). **g**, 3D surface rendering of the downstream end of a cell showed a high density of Cav-1 where F-actin accumulates. **h**, Representative image of merged Cav-1, ZO-1 and nuclear staining used for Cav-1 cluster analysis. The cluster index was determined in subcellular segments, namely upstream, mid and downstream regions, and then plotted as a bar graph, demonstrating larger clusters at the downstream end. Shown are means + SD from n = 20 cells. Data was analyzed by one-way ANOVA and post hoc Tukey’s multiple comparison test; \*\*\*\**P* < 0.0001 (Upstream vs. Downstream; Mid vs. Downstream).

To further investigate the connection between flow and caveolae distribution, we cultured human aortic EC (HAEC) monolayers on y-shaped chambers and exposed them to unidirectional laminar flow for 48 h. HAECs in the high-flow region (∼20 dynes/cm^2^) collectively aligned with the flow direction and were morphologically elongated, with a three-fold increased cell aspect ratio compared to HAECs in the low-flow region (∼5 dynes/cm^2^) (Fig. S1b). Again, Cavin-1 and Caveolin-1 showed high concentration at the downstream end of HAECs exposed to high flow (Fig. 1b; Fig. S1d).

We examined the broader cell surface asymmetries of flow-aligned HAECs by atomic force microscopy (Fig. S1e). The Young’s modulus, as calculated from the force curves, showed that the cell’s downstream end was considerably stiffer, with the average Young’s modulus (7.0 kPa) being higher compared to that of the upstream end (5.3 kPa) (Fig. S1f). Furthermore, Laurdan dye imaging revealed asymmetry of the membrane fluidity across the ECs, where the downstream end typically exhibited a higher generalized polarization (GP) and higher membrane rigidity compared to the upstream end (Fig. 1c). Membrane fluidity is a result of the distribution and composition of lipids within the bilayer. The fluorescent probe BODIPY FL C_5_-Ganglioside G_M1_ was strongly enriched at the downstream end of flow-aligned HAECs (Fig. 1d), suggesting the presence of liquid-ordered or lipid raft domains in regions with enriched Caveolin-1^32–34^.

F-actin also displayed densely webbed clusters at the downstream end of flow-aligned HAECs (Fig. 1e). Importantly, these downstream F-actin clusters correlated with high density Caveolin-1 staining (Fig. 1f). This was further highlighted for Caveolin-1 and F-actin by 3D surface rendering (Fig. 1g). Significantly, the Caveolin-1 cluster size was increased ca. six-fold within these polarized downstream areas (Fig. 1h). In summary, exposure to sustained high laminar flow resulted in the formation of domains at the downstream end of cells, characterized by higher membrane rigidity, and the accumulation of raft-type lipids, F-actin aggregation and caveolae.

### Activation of eNOS and Ca^2+^ entry occur at the downstream end

Given the known functional connections between caveolae and eNOS^20, 21, 35, 36^, we hypothesized that polarization of Caveolin-1-rich membrane domains led to localized eNOS activation. Indeed, eNOS phosphorylated on serine 1177 (p-eNOS), an active form of eNOS, was found predominantly concentrated at the downstream end of flow-aligned HAECs, with its staining mostly overlapping with that of the polarized Caveolin-1 clusters (Fig. 2a). Thus, Caveolin-1 polarization correlates with localized activation of eNOS at the downstream end of ECs exposed to sustained flow.

**Fig. 2:**
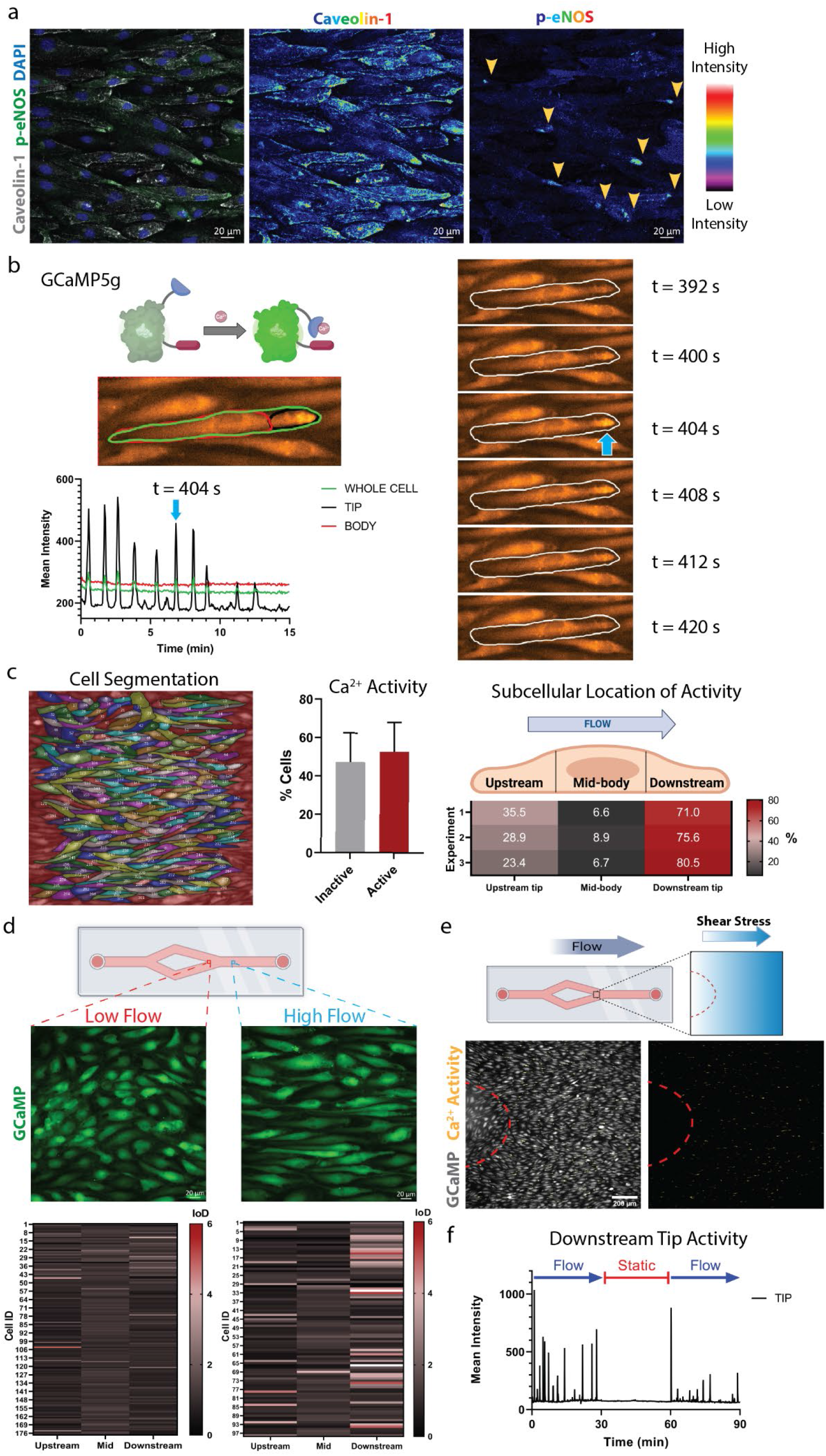
eNOS phosphorylation and Ca^2+^ oscillations occur at the downstream end in the presence of high laminar flow. For (a-c), confluent monolayers of HAECs were exposed to laminar shear stress (20 dynes/cm^2^) for 48 h. **a**, Representative image of flow-aligned HAECs stained for Cav-1 and eNOS phosphorylated on S1177 (p-eNOS). Individual channel images, displayed in Rainbow Lookup Table (LUT), highlight the accumulation of signal for both Cav-1 and p- eNOS at the downstream end (arrowheads). **b**, Imaging of Ca^2+^ activity in live HAECs transfected to express GCaMP. Fluorescence intensity over time is plotted for 15 min for one full-length cell using 3 defined regions of interest: “whole cell” (green), “tip” (black) and cell “body” (red). Note that Ca^2+^ oscillations are observed exclusively at the tip; blue arrow indicating one Ca^2+^ peak. Corresponding time sequence of the cell is displayed for indicated timepoints and blue arrow indicates the same Ca^2+^ peak on the fluorescence trace. **c**, All cells within an imaging field of view were outlined and ID’ed as shown in the representative image on the left. The GCaMP signal was extracted over 30 min from 732 cells (n = 3 independent experiments) and then analyzed. Approximately 50 % of the cells had Ca^2+^ activity (index of dispersion, IoD, greater than 2). Active cells were further segmented into 3 equal-length segments for the upstream, mid-body and downstream subcellular regions. Of the active cells, over 70% had Ca^2+^ activity restricted to the downstream end. **d,** To model low-flow and high-flow areas, HAECs were seeded on y- shaped slides and exposed to unidirectional laminar flow for 48 h prior to imaging. Ca^2+^ activity in low-flow (∼5 dynes/cm^2^) and high-flow regions (∼20 dynes/cm^2^) was imaged in GCaMP transfected HAECs. Cell segmentation and extraction of the fluorescence intensity over time showed enhanced activity at the downstream tip in the high-flow region. IoD plots show data from 117 low-flow and 98 high-flow cells across n = 3 biological replicates. **e,** Region of the y-shaped slide, where flow converges, experience increasing shear stress levels. Cells in the high-flow region were morphologically aligned and exhibited more Ca^2+^ spikes (in yellow) compared to the unaligned cells in the low- flow area. **f,** Confluent monolayers of GCaMP transfected HAECs were exposed to laminar shear stress (20 dynes/cm^2^) for 48 h prior to live cell imaging. Representative GCaMP intensity trace showing 90 min of Ca^2+^ activity at the downstream tip in the presence of flow (30 min; 20 dynes/cm^2^), static (30 min; 0 dynes/cm^2^) and re-flow (30 min; 20 dynes/cm^2^) conditions.

Phosphorylation of eNOS on serine 1177 is frequently Ca^2+^ dependent^37, 38^. To examine changes in intracellular free Ca^2+^, we transfected HAECs with plasmids encoding the Ca^2+^ reporter GCaMP and exposed them to high laminar flow (Fig. 2b; Movie 1). Segmentation analysis of individual cells revealed that oscillatory Ca^2+^ influx events were restricted to the downstream end, previously identified as enriched for Caveolin-1 and p-eNOS (Fig. 2b; Movies 2 & 3). To quantify the prevalence of Ca^2+^ activity across the monolayer, full-length cells were segmented and ‘active’ cells were identified as having an index of dispersion (IoD) greater than 2. Approximately 50% of the cells showed Ca^2+^ transients over the 30 min of imaging (Fig. 2c), indicating that oscillatory Ca^2+^ influx events were a sustained response of ECs under high laminar flow. Segmentation analysis of these active cells revealed that transients were restricted to the downstream end in over 70% of the active cells (Fig. 2c; Fig. S2a).

### Polarized signaling activity requires high laminar flow

We next compared Ca^2+^ activity in low flow versus high flow HAECs on y-slides and observed that elevated shear stress correlated with increased Ca^2+^ oscillations. In the low-flow region, cells were less elongated and exhibited less Ca^2+^ oscillations, which were not preferentially observed within specific subcellular domains (Fig. 2d; Movies 4 and 5). In regions of increasing shear stress, cell morphology changed from cobblestone to elongated, and we observed concomitant enhanced Ca^2+^ activity with increasingly preferential localization to the downstream end of the elongated cells (Fig. 2e; Fig. S2b; Movie 6). In fact, there was a positive correlation between cell aspect ratio and Ca^2+^ oscillations at the downstream tip (Fig. S2c), suggesting that flow-induced elongation is required for increased Ca^2+^ signaling. To confirm the requirement of shear stress for the Ca^2+^ signaling, we imaged GCaMP-HAECs over the time course of flow, static and re- flow conditions to reveal that the oscillatory Ca^2+^ activity was observed only in the presence of laminar flow (Fig. 2f; Fig. S2d; Movie 7).

To investigate whether cell body elongation was sufficient for these flow-induced phenotypes, we elongated HAECs by culturing them on line-patterned slides to achieve cell aspect ratio similar to those seen with flow, but in the absence of flow (Fig. 3a). Caveolin-1 showed asymmetric distribution at one end of static, elongated cells (Fig. 3b). However, in contrast with flow-aligned and elongated HAECs, static elongated ECs accumulated Caveolin-1 on either the left or right side of the cell, resulting in a random pattern. Importantly, polarized p-eNOS was not observed in static elongated ECs (Fig. 3c). Accordingly, they did not display Ca^2+^ oscillations as measured by live cell imaging of GCaMP-HAECs (Fig. 3d; Movie 8). Unlike HAECs exposed to high flow, HAECs exposed to low flow and static elongated HAECs did not show asymmetric activation of p-eNOS (Fig. 3e). Thus, while cell elongation was sufficient to trigger Caveolin-1 clustering at either end of HAECs, the sustained presence of high laminar flow was required for coordinated polarization of Caveolin-1 within the population, and the formation of functional signaling domains.

**Fig. 3:**
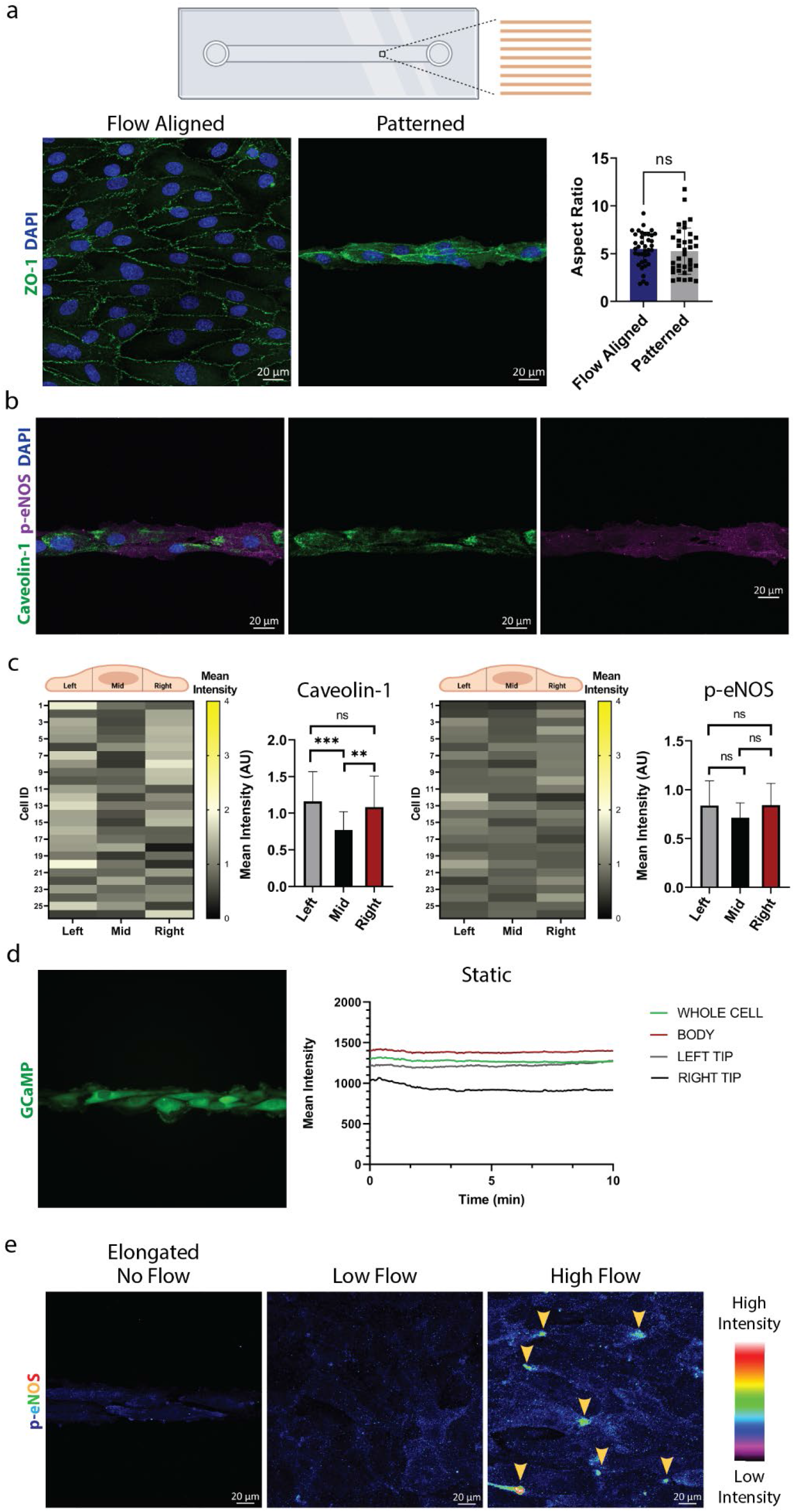
High laminar flow is required for localized signaling activity. **a,** HAECs were seeded on either non-patterned chambers and flow aligned (20 dynes/cm^2^) (left panel), or seeded on line-patterned chambers and cultured statically. After 48 h, cells were analyzed and aspect ratio calculated as for Supplemental Fig. 1b. Shown are means ± SD. ns, not significant by two-tailed, unpaired *t* test. **b, c,** HAECs were elongated statically on the line-patterned chamber and stained for Caveolin-1, p- eNOS and DAPI. Representative images of the staining are shown in (b), with segmentation analysis from 26 cells shown in (c). Bar charts show intensity means ± SD data and analyzed by one-way ANOVA with post hoc Tukey’s multiple comparisons test. ns, not significant, \*\**P* < 0.01, \*\*\**P* < 0.001. **d,** Ca^2+^ activity was recorded for GCaMP- expressing HAECs that were cultured statically on line-patterned chambers. Representative live cell recording of intensity trace for one cell over 10 min showed a complete lack of localized activity. **e,** HAECs were cultured statically on the line-patterned chamber or y-shaped slide for 48 h. Representative images of p-eNOS staining using equivalent imaging conditions to compare the signal intensity across conditions. The p- eNOS signal is displayed using false color Rainbow Lookup Table (LUT) to highlight the clustered regions of staining in cells under high flow.

### Localized Ca^2+^ entry occurs via Trpv4/Caveolin-1 association at the downstream end

We postulated that Ca^2+^ entry from the extracellular space via plasma membrane channels was required for Ca^2+^ activity. Consistent with this, the addition of the Ca^2+^ chelator EGTA blunted these Ca^2+^ oscillations (Fig. 4a; Fig. S3a). The Trpv4 ion channel is implicated in endothelial Ca^2+^ ‘sparklet’ activity in mouse arteries^39–42^. To test the role of Trpv4 in Ca^2+^ activity, we added the Trpv4 antagonist GSK205 during imaging of GCaMP-HAECs under flow and found that this treatment suppressed Ca^2+^ activity (Fig. 4b; Fig. S3b). This indicated that Trpv4 is required for oscillatory Ca^2+^ entry at polarized Caveolin-1-rich regions where active eNOS is observed in flow-exposed elongated ECs.

**Fig. 4:**
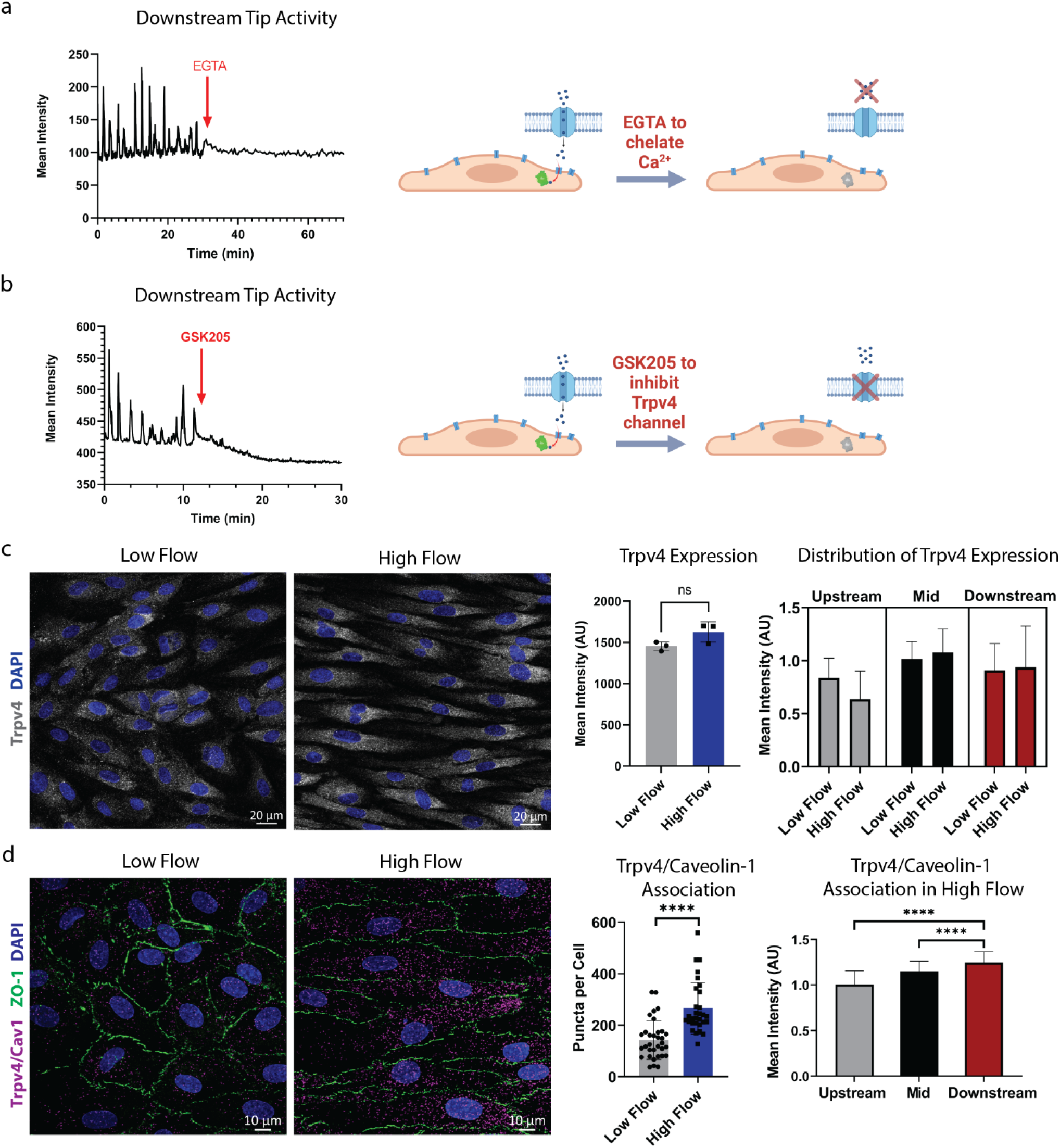
Localized Ca^2+^ entry requires Trpv4 channel activity and occurs in areas of Trpv4/Caveolin-1 association. For (a-b), confluent monolayers of GCaMP transfected HAECs were exposed to laminar shear stress (20 dynes/cm^2^) for 48 h prior to live cell imaging. **a,** Representative GCaMP intensity trace showing Ca^2+^ activity at the downstream tip after the addition of EGTA (1.6 µM) to chelate calcium ions in the culture media. **b,** Representative GCaMP intensity trace showing Ca^2+^ activity at the downstream tip after the addition of the Trpv4 antagonist GSK205 (20 µM). For (c-d), HAECs were seeded on y-shaped slides and exposed to unidirectional laminar flow for 48 h prior to staining. Immunofluorescence was compared for cells in low-flow (∼5 dynes/cm^2^) and high-flow regions (∼20 dynes/cm^2^). **c,** Trpv4 protein staining showed no difference for low flow versus high flow regions (representative images from n = 3 biological replicates; statistics calculated by two-tailed, unpaired *t* test to show no significance, ns, between regions). Quantifying the subcellular distribution of expression indicates that Trpv4 was not polarized under flow. Shown are means + SD from 44 low-flow and 57 high-flow cells. **d,** Representative images of ZO-1 staining (green) and proximity ligation assay (PLA) to detect Trpv4 and Caveolin-1 association (magenta puncta) in low-flow and high-flow regions. Cells in the high-flow region showed a significant increase in PLA puncta per cell compared to cells in the low-flow region. Shown are puncta/cell as well as mean ± SD and statistics calculated using unpaired, two-tailed *t* test; \*\*\*\**P* < 0.0001. Additional segmentation analysis showed that Trpv4 and Caveolin-1 PLA puncta preferentially occurred in the downstream end. 111 cells were analyzed. Data was analyzed by one-way ANOVA and post hoc Tukey’s multiple comparisons test; \*\*\*\**P* < 0.0001 (Upstream vs. Downstream; Mid vs. Downstream).

Next, we investigated how polarized Trpv4-dependent Ca^2+^ entry was produced in response to flow conditioning of HAECs. Trpv4 mRNA and protein levels were indistinguishable in HAECs irrespective of the presence of laminar flow, whereas known flow-responsive genes KLF2 and KLF4, and eNOS protein showed significant upregulation in response to flow (Fig. S3c, S3d). Importantly, unlike other proteins such as Caveolin-1, Cavin-1, or F-actin, Trpv4 distribution did not display any apparent polarization (Fig. 4c). Trpv4 can interact with Caveolin-1, and this association leads to its activation in lung endothelium^39^. Here, we used proximity ligation assay (PLA) to determine if such an association also occurred in flow-conditioned HAECs. We observed specific *in situ* PLA spots when combining antibodies to Caveolin-1 and Trpv4, or Cavin- 1, but not Histone H3 or with individual antibodies alone (Fig. 4d; Fig. S3e). Significantly, Trpv4/Caveolin-1 PLA spots were strongly enhanced in HAECs exposed to high flow and also clearly accumulated at the downstream end of the elongated cells. Cells that are not morphologically elongated in the low-flow region did not show a difference in distribution of PLA spots (Fig. S3f). Combined, these data indicated that Trpv4 associates with Caveolin-1 clusters in the presence of high flow within downstream regions of the cell where oscillatory Trpv4-mediated Ca^2+^ activity occurs.

### Disruption of polarized caveolae abolishes localized Ca^2+^ activity

Cholesterol affects Caveolin-1 clustering and functions of caveolae, and sequestration of cholesterol using methyl-β-cyclodextrin (MβCD) disrupts caveolae function^43–45^. We treated flow-aligned HAECs with MβCD for 30 min (Fig. 5a) to deplete the accessible pool of cholesterol (as measured with the probe ALO-D4^46^) (Fig. S4a) and disrupt flow-induced plasma membrane polarization of Caveolin-1 (Fig. 5b). Functionally, this treatment also reduced the levels of intracellular NO (Fig. 5c), further supporting that Caveolin-1 polarization directly affects endothelial signaling. Indeed, depletion of plasma membrane cholesterol also resulted in a loss of localized Ca^2+^ activity under laminar flow (Fig. 5d-f). Intriguingly, treatment with the Trpv4-specific agonist GSK1016790A (GSK101; 10 nM) led to Ca^2+^ activity at the ends of elongated ECs exposed to MβCD (Fig. 5d-f; Movie 9), indicating that Trpv4 channels continued to be activatable. Considering that Trpv4 channel expression was observed across the entire cell surface (Fig. 4c), we conjectured that Ca^2+^ entry through the channel could be modulated by the local membrane environment. In line with this hypothesis, the addition of a higher dose of GSK101 (1 µM) resulted in Ca^2+^ entry throughout the entire cell body (Fig. S4b) suggesting that the Trpv4 channel open probability was greater at the ends of flow-elongated endothelial cells.

**Fig. 5:**
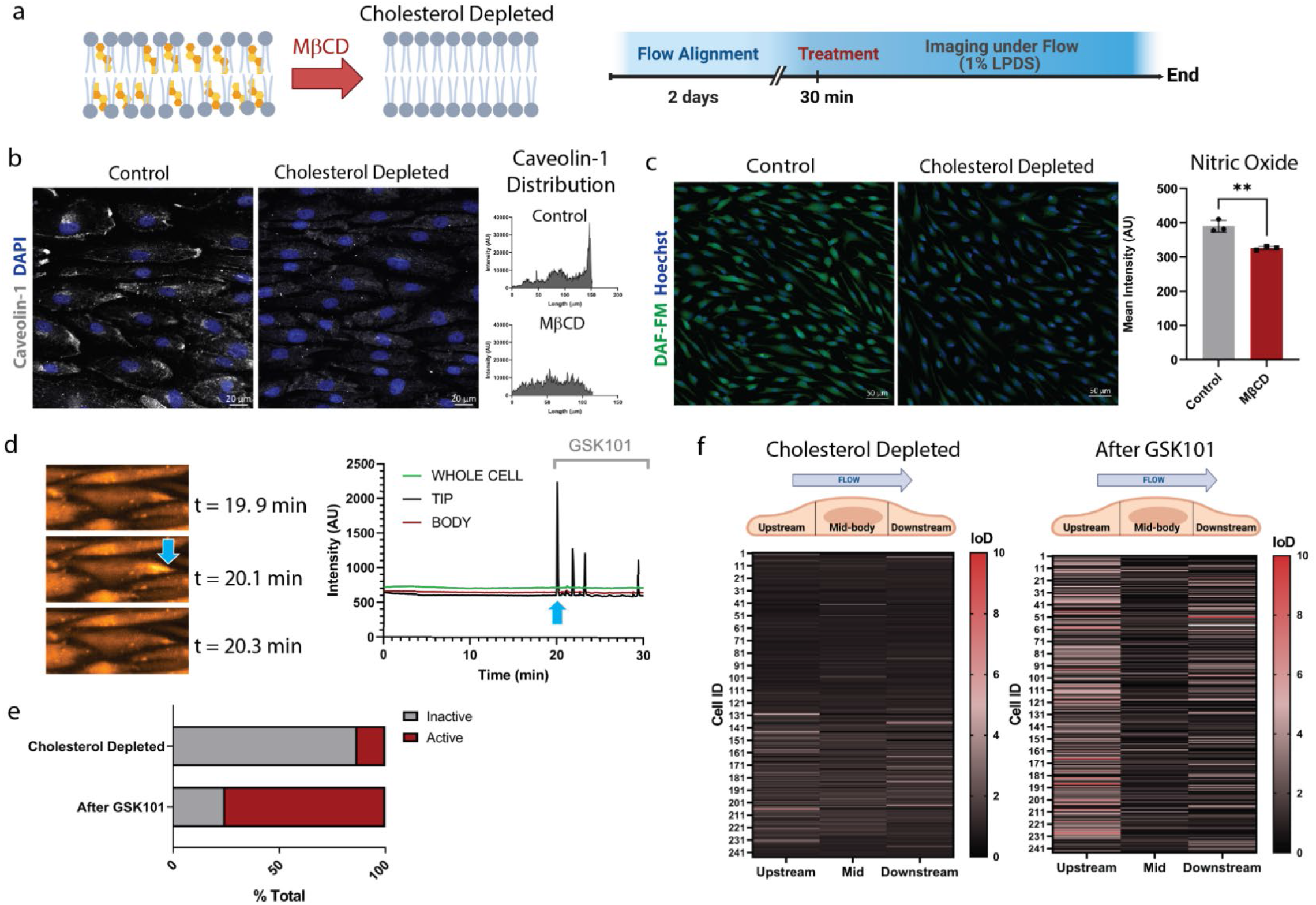
Cholesterol depletion abolishes polarized signaling. **a,** MβCD was used to deplete plasma membrane cholesterol. **b,** Flow-aligned HAECs were treated with MβCD for 30 min then fixed and stained for Caveolin-1 and DAPI. MβCD treatment abolished Caveolin-1 polarization as clearly shown by intensity plots of representative cells from the control and MβCD treated groups. **c,** NO production was visualized via DAF-FM loaded flow-aligned monolayers of control and MβCD treated groups. Shown are mean DAF-FM fluorescence intensities ± SD for n = 3 biological replicates and statistics calculated using two-tailed, unpaired *t* test; \*\**P* = 0.0044. **d,** GCaMP imaging of the cholesterol-depleted cells under flow (20 dynes/cm^2^) for 20 min showed lack of Ca^2+^ activity. Displayed are time-dependent images of a representative cell and the corresponding intensity trace for the whole cell and indicated segments. At t = 20 min (blue arrow), the Trpv4 agonist GSK1016709A (GSK101, 10 nM) was added to the flowing culture media. This led to an immediate Ca^2+^ burst as seen in the image at 20.1 min. **e,** Overall, only 13% of the cells depleted for cholesterol were active in the initial 20 min of imaging. The number of active cells increased to 75% following the addition of GSK101. **f,** IoD heatmaps show Ca^2+^ activity following cholesterol depletion and subsequent GSK101 addition for n = 244 cells.

### Polarized Trpv4 activity limits inflammation

We next investigated whether flow-induced Trpv4 signaling activity promotes an anti-inflammatory response. In cells exposed to continued laminar flow, GSK205-mediated inhibition of Trpv4 activity (Fig. 6a) resulted in reduced NO production (Fig. 6b) and increased reactive oxygen species (ROS) generation (Fig. 6c), whereas GSK101- mediated activation of Trpv4 activity enhanced NO production (Fig. S5a). Moreover, GSK205 treatment also led to increased surface expression of the pro-inflammatory adhesion molecules ICAM-1 and E-Selectin (Fig. 6d), alongside enhanced nuclear localization of the NF-κB p65 subunit (Fig. 6e). We confirmed that Caveolin-1 rich domains remained polarized during Trpv4 inhibition (Fig. S5b). Thus, flow-induced Trpv4 activation promotes an anti-inflammatory phenotype.

**Fig. 6:**
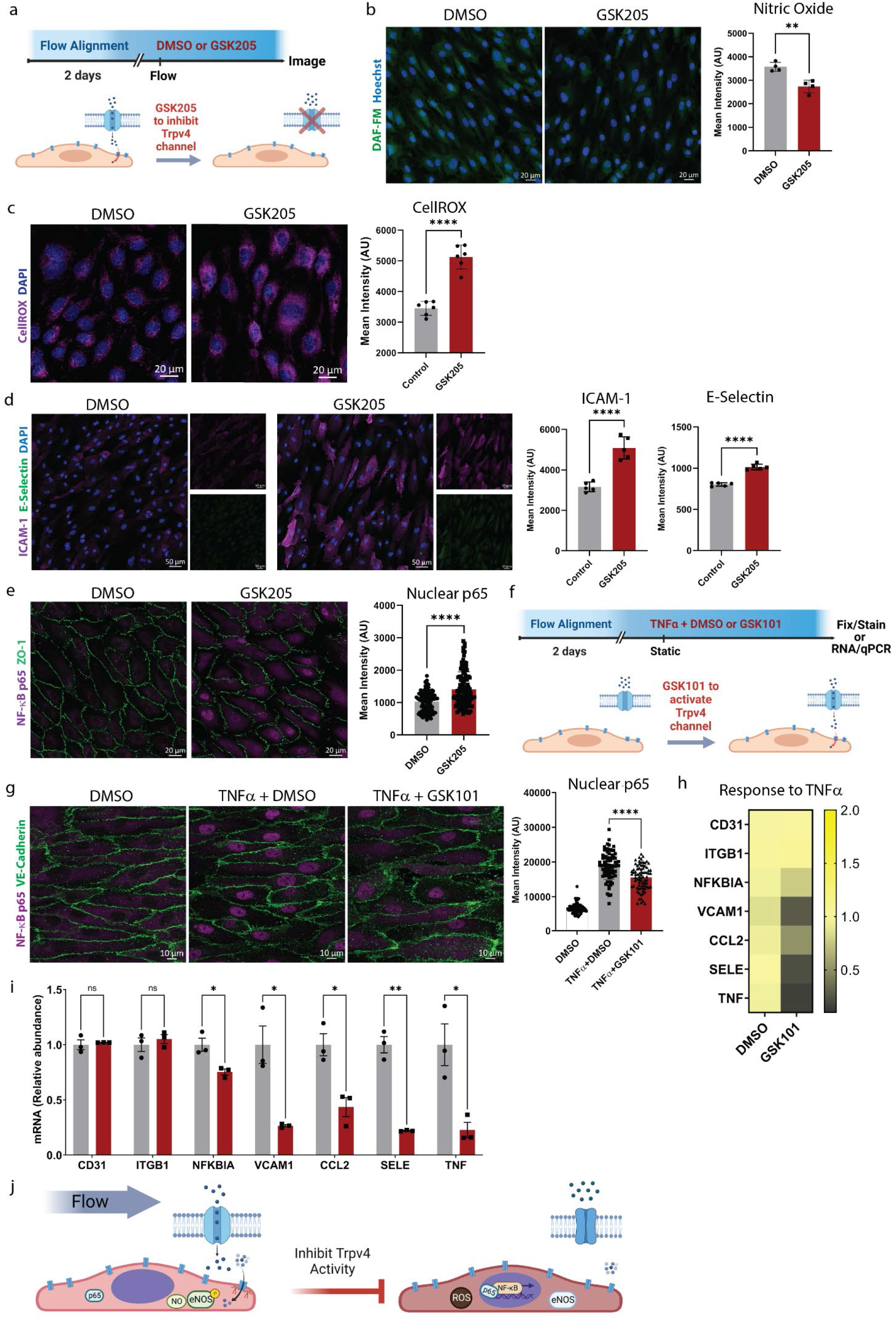
Trpv4 activity limits the inflammatory response in HAECs. **a,** HAEC monolayers were flow aligned for 48 h followed by the addition of DMSO (control) or Trpv4 antagonist GSK205 (20 µM) in the presence of laminar flow (20 dynes/cm^2^). **b,** Quantification of NO production in live cells by DAF-FM imaging after 2 h of treatment. Shown are representative images and mean intensities ± SD from n = 3 biological replicates and statistics calculated using two-tailed, unpaired *t* test; \*\**P* = 0.0023. **c,** After 2 h treatment, control and GSK205 treated monolayers were incubated with CellROX probe and imaged to quantify reactive oxygen species. Shown are representative images and mean intensities ± SD from n = 6 biological replicates and statistics calculated using two-tailed, unpaired *t* test; \*\*\*\**P* < 0.0001. **d,** Control and GSK205 treated monolayers were fixed after 4 h and stained for ICAM-1 and E-Selectin. Shown are representative images and mean intensities ± SD from n = 5 biological replicates and statistics calculated using two-tailed, unpaired *t* test; \*\*\*\**P* < 0.0001 for both graphs. **e,** Control and GSK205 treated monolayers were fixed after 2 h and stained for DAPI, junctional marker ZO-1 and the p65 subunit of NF-κB. Nuclear expression of NF-κB p65 was quantified by mean fluorescence intensity after nuclear mask application. Shown is mean intensity from n = 115 cells analyzed for each condition for n = 3 biological replicates and statistics calculated using two-tailed, unpaired *t* test; \*\*\*\**P* < 0.0001. **f,** Experimental design for static TNFα treatment (10 ng/mL; 30 min) in the presence of GSK1016790A (GSK101, 10 nM) or DMSO. **g,** Following the treatment, monolayers were fixed and stained for DAPI, junctional marker VE-Cadherin and NF-κB p65. Graph represents mean fluorescence intensity in the nucleus from n = 79 cells per group and statistics calculated using one-way ANOVA with post hoc Tukey’s multiple comparisons test; \*\*\*\**P* < 0.0001. **h,** Gene expression was measured by qPCR for TNFα treated monolayers in the presence of DMSO or GSK1016790A (GSK101). Gene expression of NFKBIA, VCAM1, CCL2, SELE and TNF is plotted as a heat map of the mean expression (n = 3 biological replicates). Note that CD31 and ITGB1 are not changed with GSK101 treatment. **i,** as in (h) with mRNA expression plotted as mean ± SEM for the inflammatory genes, CD31 and ITGB1. \**P* < 0.05, \*\**P* < 0.01 by two-tailed, unpaired *t* test. **j,** Graphical model describing how laminar flow supports localized Trpv4 activation by polarized Caveolin-1 domains, which leads to Ca^2+^ activity, eNOS activation, NO production and inhibition of NF-κB mediated transcription.

To further determine whether induction of Trpv4 activity could enhance an anti-inflammatory response in ECs, we used flow-aligned HAECs in static conditions to study the effect of GSK101 and Tumor Necrosis Factor alpha (TNFα) (Fig. 6f). Caveolin-1-rich domains remained polarized for at least 30 min after removal of flow (Fig. S5c). Following a 30 min treatment with TNFα we observed nuclear translocation of NF-κB p65 (Fig. 6g). However, this response was attenuated upon activation of Trpv4 by treatment with GSK101 (10 nM) (Fig. 6g). GSK101 significantly reduced the TNFα-mediated upregulation of pro-inflammatory gene expression including Nuclear Factor-kappa-B-Inhibitor alpha (NFKBIA), Vascular Cell Adhesion Molecule 1 (VCAM1), C-C Motif Chemokine Ligand 2 (CCL2), Selectin E (SELE), and TNF (Fig. 6h and Fig. 6i). Changes to ICAM-1 protein expression levels were not observed within this short inflammatory TNFα stimulation (Fig. S5d). However, the dampening of TNFα-stimulated ROS production with GSK101 co-stimulation (Fig. S5e) indicated that ectopic Trpv4 activation suppresses the inflammatory TNFα response in HAECs.

## Discussion

Our work shows that flow-induced elongation preferentially polarizes Caveolin-1 rich domains to the downstream end of arterial ECs. This polarization in the presence of laminar flow activates Trpv4 and eNOS in a spatially restricted manner to suppress inflammatory pathways (Fig. 6j). We provide clear evidence that Caveolin-1 clusters at the downstream end of flow-aligned aortic ECs *in vivo* and *in vitro*. We reveal that these clusters define a previously undescribed mechanosensitive domain that enables focal Ca^2+^ entry for eNOS activation and inhibition of inflammatory signaling. These domains were associated with a distinct pattern of proteins, lipids and consequently biophysical properties rendering the downstream luminal surface of flow-aligned HAECs significantly different to other parts of the cell. It is yet unclear how elongation and laminar shear stress induce this local modulation of the plasma membrane, with both cytoskeletal and membrane lipids possible mediators^47–50^.

Caveolin-1 and caveolae have been long recognized as important regulators of arterial organization in response to shear stress, in particular with respect to the regulation of vascular tone^51, 52^. Our data shows that aortic Caveolin-1 also plays an important role in promoting endothelial cell resilience^53^ by dampening inflammatory gene expression. Caveolin-1 clustering can lead to caveolar rosettes to provide additional membrane in response to elevation of surface tension^54, 55^. Future investigation of how Caveolin-1 clustering at the downstream end of flow-aligned HAECs affects the organization and dynamic redistribution of caveolae and membrane lipid composition will thus be of particular interest. The contribution of the actin cytoskeleton in stabilizing these domains also warrants further investigation.

Most importantly, these membrane domains enabled locally restricted signaling as shown by focal Ca^2+^ entry via Trpv4 channels and phosphorylation of eNOS at the downstream end of aortic ECs in the presence of high laminar flow. The mechanosensitivity of these domains was clearly shown by cycling the applied flow on and off to reveal that Ca^2+^ oscillations were only sustained when shear stress was present. Furthermore, since all imaging experiments were performed after 48 h of flow exposure, these Ca^2+^ oscillations were associated with the continuous presence of flow, in contrast to previous observations of Ca^2+^ spikes at the onset of shear stress^56–58^.

While Trpv4 has been identified as a mechanosensitive ion channel^59–61^, the modulation of its activity in the plasma membrane is less clear. The physical association of Trpv4 with Caveolin-1 at the downstream end of the cell indicates that this activation mechanism plays an important role in restricting Ca^2+^ entry to this region of the cell. Since inhibiting the Trpv4-mediated Ca^2+^ influx resulted in reduced NO production and an increase in both ROS and inflammation, our data suggests that endothelial cell resilience is clearly dependent on the presence of these polarized plasma membrane domains. While endothelial Trpv4 activity has been shown to promote vasodilation in arteries^31, 62–64^, its role in vascular inflammation has not been as clearly defined^65, 66^. In fact, there is conflicting data regarding the role of Trpv4 activity in the inflammatory cascade^67, 68^. Here, we provide evidence that Trpv4 activity under laminar flow is anti-inflammatory in aortic ECs. It led to eNOS activation and the production of NO, both indicators of anti-inflammatory signaling^69, 70^, and ultimately contributed to dampening of inflammatory gene expression. Therefore, this localized Trpv4 activation is a previously unrealized athero-protective mechanism of laminar flow and eNOS activation. Future research will establish whether additional protective functions of laminar flow are mediated by polarized Caveolin-1/Trpv4 domains.

Additional studies are required to determine whether there exists a critical shear stress level or ‘set point’^71^ to initiate and sustain the localized signaling activity. While our studies were limited to laminar shear stress, future experiments will also determine whether there are differences in the compartmentalization of signaling activity in the presence of disturbed/oscillatory flow. In conclusion, we reveal that the anterior-posterior endothelial cell polarity established in response to laminar flow results in mechanosensitive signaling domains that both promote cell resilience and suppress inflammation.

## Materials and Methods

### Cell Culture and Shear Stress

Either primary HAECs (Cell Applications #S304-05a) or immortalized TeloHAECs (ATCC #CRL-4052) were used. Primary HAECs were used from P4-P12, and TeloHAECs were used up to P20. For all cell culture experiments, MCDB-131 complete media (VEC Technologies #MCDB-131 Complete) was supplemented with 10% FBS (Omega USDA certified FBS #FB-11). For plating cells on cell culture dishes, 0.1% gelatin (Stemcell #07903) coating was first applied. Cells were cultured in a 37 °C incubator with 5% CO_2_.

For application of shear stress, endothelial cells were seeded in ibidi µ-Slide 0.4 Luer ibiTreat (ibidi #80176) or y-shaped ibiTreat chambers (ibidi #80126). Unidirectional flow was applied to confluent monolayers using the ibidi pump system (ibidi #10902). For experiments requiring access to the cells, including atomic force microscopy, cell monolayers were exposed to shear stress in glass bottom 6-well plates (Cellvis #P06- 1.5H-N) or 35mm glass-bottom FluoroDishes (WPI #FD35-100) on an orbital shaker (Benchmark Scientific #BT302). A rotation speed of 130 rpm was applied to achieve endothelial cell alignment on the periphery of the well where the flow is unidirectional, and cells are unaligned in the center of the well where flow is multi-directional^72, 73^. For cell alignment in the absence of flow, cells were plated on customized ibidi µ-slides with multi-cell micropattern (ibidi GmbH).

### Atomic Force Microscopy (AFM)

Surface measurements were performed on HAECs using JPK NanoWizard 4a AFM and Bruker PFQNM-Live Cell probe with a spherical tip (Bruker AFM Probes #PFQNM-LC) and tip length 17 µm and tip radius 65 nm. The spring constant of the cantilever was determined by manufacturer and inputted into JPK acquisition software. Imaging was performed using QI™ mode for quantitative imaging to capture the entire cell and plot the height profile. Force mapping was then performed at the two ends of the cell as user defined 15 x 32 pixel regions with tapping frequency of 10 Hz and probe extend speed of 3 µm/sec and 0.6 nN setpoint applied. To quantify the Young’s modulus, JPK Data Processing analysis software was used to fit individual force curves using Hertz model. For identification of the cell surface properties, the indentation depth was set to less than 100 nm for each force curve to keep within 0.1% strain.

### Live Cell Fluorescence Imaging

#### Membrane Fluidity

For visualization of cell plasma membrane fluidity, Laurdan dye (6- Dodecanoyl-2-Dimethylaminonaphthalene) (Invitrogen #D250) was applied at 10 μM to cell monolayers and incubated 30 min at 37°C then washed and imaged in 1X HBSS. For imaging, Zeiss LSM 880 with Chameleon 2-photon laser with Plan-Apochromat 20x/0.8 M27 objective and GaAsP PMT array detector were used. Images were collected using two-photon excitation set to 770 nm, and two-channel emission detection was set for 400 – 460 nm & 470 – 530 nm. The acquisition of generalized polarization (GP) images was performed using the ImageJ (National Institutes of Health) software and ImageJ macro GPCalc ratiometric quantification and visualization^74^.

#### Lipid Raft Imaging

Live cell imaging was performed using green fluorescent BODIPY™ FL C_5_-Ganglioside G_M1_ (Invitrogen #B13950). Cell monolayers were treated with 50 µM BODIPY™ FL C_5_-Ganglioside G_M1_ for 20 minutes at 37°C with Hoescht 33342 (AAT Bioquest #17535).

#### Calcium Imaging

HAECs were transfected with GCaMP plasmid (pPB_CAG_GCamp5g) with PiggyBac construct (PB_Vector) (gift from Dr. Roy Wollman, UCLA) using Lipofectamine 3000 transfection reagent (Invitrogen #L3000001) in Opti-MEM media (Gibco #31985062) overnight. Cells were then selected for GCaMP expression with Blasticidin (Gibco #A1113903). GCaMP expressing HAECs were plated on Y-shaped chamber slides (ibidi #80126) and after reaching confluence, slides were connected to ibidi pump system for flow conditioning. After 48 h of unidirectional flow, the y-slide was connected to yellow/green perfusion set (ibidi #10963) modified with non-permeable tubing containing 13 mL of conditioned MCDB-131 (VEC Technologies #MCDB-131 WOFBS) with 10% FBS (Omega USDA certified FBS #FB-11) for live cell imaging. All steps were completed with minimal light exposure. Fluorescence images were acquired using Zeiss Observer Z1 with Colibri 7 light source, CMOS camera (Photometrics Prime 95B) and ZEN Blue software. Images were acquired once every 3 s for total imaging time. For observation of changes in calcium activity, chemicals were added to one of the ibidi syringe reservoirs when air pressure was not active. Chemicals used include ethylene glycol tetra-acetic acid (EGTA,1.6 mM; Fisher Scientific #NC1280093), GSK1016790A (10 nM or 1 µM; Sigma-Aldrich #530533), GSK205 (20 µM; MedChemExpress #HY- 120691A).

#### Nitric Oxide Imaging

For quantification of nitric oxide production, cells were treated with 5 µM DAF-FM (Invitrogen #D23844) for 20 min at 37°C with Hoechst 33342 (AAT Bioquest #17535).

#### Reactive Oxygen Species Imaging

For oxidative stress detection, cell monolayers were treated with 5 µM CellROX Deep Red Reagent (Invitrogen #C10422) or 5 μM CM- H_2_DCFDA (Molecular Probes, C6827) for 20 min at 37°C with or without Hoechst 33342 (AAT Bioquest #17535).

#### Membrane Cholesterol Imaging

Visualization of the accessible pool of cholesterol was performed after 30 min treatment with 10 mM methyl-beta-cyclodextrin (MβCB) in 1% lipoprotein-deficient serum (LPDS, Sigma #S5519) or 1% LPDS control, followed by flow for 2 hours in complete media. HAECs were then stained with 20 μg/mL ALOD4-488 (gift from Peter Tontonoz, UCLA) for 10 min at room temperature followed by 4% paraformaldehyde (PFA, Sigma-Aldrich #322415) fixation and application of 4,6- Diamidino-2-phenylindole, dihydrochloride (DAPI) nuclear stain (AAT Bioquest #17507).

After respective dye incubation, cell monolayers were washed with 1X HBSS then imaged in culture media or 1X HBSS using Zeiss Colibri LED plus CMOS camera (Photometrics 95B) detection or LSM900 with Airyscan2 GaAsP-PMT detector for gentle confocal imaging.

### Proximity Ligation Assay

The Duolink proximity ligation assay (PLA) was performed according to the manufacturer’s protocol (Sigma Aldrich #DUO92008). Briefly, HAECs were grown on Y-shaped chamber slides (ibidi #80126) and after reaching confluence, slides were connected to ibidi pump system for flow (20 dynes/cm^2^, unidirectional laminar flow) for 48 h. Then, cells were fixed with 4% PFA for 5 min at room temperature. After washing with 1X PBS, the cells were permeabilized with 0.05% Triton X-100 (Fisher Scientific #A16046-0F) in 1X PBS for 5 min, blocked in Duolink blocking solution for 60 min at 37°C and then incubated with primary antibody (α-Trpv4, LSBio #LS-C401108 and α-Caveolin- 1, R&D #AF5736; α-Trpv4 or α-Caveolin-1 only or α-Caveolin-1 and α-nuclear HistoneH3 Abcam #ab8898, as negative controls; α-Caveolin-1 and α-Cavin-1, Abcam #ab48824, as a positive control) diluted in Duolink Antibody Diluent overnight at 4°C. After washing, the cells were incubated with PLA probes including PLUS-Goat (#DUO92003), MINUS- Rabbit (#DUO92005) at 37°C for 1 h. After ligation, amplification, and final washes, the cells were counter-stained with ZO-1 conjugated with AF488 (Invitrogen #MA3-39100- A488) and DAPI (AAT Bioquest #17507). Images were obtained using Zeiss LSM900 with Airyscan2 GaAsP-PMT detector.

### Animals

Male and female wildtype C57BL/6J mice were purchased from The Jackson Laboratory (Strain #000664). All mice were fed a chow diet and water ad libitum under a 12-hour light/dark cycle. All mouse experiments were approved by the UCLA Division of Laboratory Animal Medicine (Protocol #ARC-2021-050-AM-002) and conformed to the NIH guidelines for the care and use of laboratory animals.

### Aorta En Face Immunostaining

Mice were perfused with ice cold 1X PBS with an incision of the right atrium to release the blood followed by perfusion with 4% PFA. The thoracic aorta was isolated and post-fixed with 0.4% PFA overnight at 4 °C. The vessel was then washed three times with 1X PBS and permeabilized by incubating with 0.3% Triton-X in 2% normal donkey serum (Jackson Immuno Research Laboratories #017-000-121) in 1X PBS for 30 min at room temperature followed by incubation with primary antibodies overnight at 4 °C with gentle agitation. After 1X PBS washes, the vessel was incubated with corresponding secondary antibodies for 2 h at room temperature followed by washing with 1X PBS three times for 10 min each. The vessel was then placed on a slide glass, cut longitudinally, and mounted in Fluoromount-G (SouthernBiotech #0100-01).

### Immunofluorescence and Confocal Imaging

For immunostaining, cell monolayers were fixed with 4% PFA for 10 min followed by multiple washes with 1X PBS. Samples were then blocked for 2 h with 10% Normal donkey serum (Jackson Immuno Research Laboratories #017-000-121) in 1X PBS. Depending on the protein targets of interest, samples were permeabilized with either 0.1% Triton X-100 (Fisher Scientific #A16046-0F) or 0.01% Digitonin (EMD Milipore #3004100). Primary antibodies were incubated overnight at 4 °C in blocking buffer and secondary antibodies applied for 2 h at room temperature. Primary antibodies used: Erg (Abcam #ab92513), Ve-cadherin (BD Pharmingen™ #550547), Caveolin-1 (Invitrogen #PA1-064), Caveolin-1 (R&D AF5736), Caveolin-1 (Santa Cruz #sc-894), Cavin-1 (Abcam #ab48824), Icam-1 (Santa Cruz #sc-107), E-selectin (Invitrogen #MA1-06506), ZO-1 (Invitrogen #MA3-39100-A488), NFkB p65 (Cell Signaling #8242S), p-eNOS^Ser1177^ (Invitrogen #PA5-35879), p-eNOS^Ser1177^ (Santa Cruz #sc-81510), Trpv4 (Alomone Labs #ACC-034 or LSBio #LS-C401108), VE-Cadherin (R&D #AF938). For F-actin staining, Alexa Fluor™ 555 Phalloidin (Invitrogen #A34055) was applied with secondary antibody incubation.

Imaging was performed on a Zeiss LSM 900 confocal microscope equipped with 405nm, 488nm, 561nm and 640nm laser lines using Plan-Apochromat objectives (10x, 20x, 40x or 63x) and Airyscan2 GaAsP-PMT detector. Identical laser intensity settings were applied to all samples being compared with equivalent Z thickness. After acquisition, a maximum intensity projection of the Z-stack was applied using ZEN Blue 3.5 software (ZEISS). Image processing and quantification of parameters was performed with ZEN Blue software, IMARIS software (Bitplane), ImageJ (NIH) or custom script (see section on *Image Analysis* for details).

### Image Analysis

For fluorescence intensity measurements, maximum intensity projections of the confocal Z-stack images were used. To quantify intensity differences with treatments for ALO-D4, DAF-FM, CellROX, ICAM-1, E-Selectin and Trpv4, the arithmetic mean intensity for each channel of interest was determined using ZEN blue software (Zeiss, Germany). For nuclear quantification of NF-κB p65 localization, ImageJ was used to create a binary mask on the DAPI stained nuclear channel. The nuclear mask was then applied to the NF-κB p65 stained channel using the ROI Manager and the mean fluorescence intensity was calculated per cell. For graphical tracing of the subcellular Ca^2+^ activity, the GCaMP intensity over time was exported using ZEN blue software using the ROI feature for each defined region of interest. Extracted data was plotted in GraphPad Prism software.

For analysis of the live cell GCaMP activity and confocal imaging, segmentation was performed by hand-drawing cell borders to input mask and raw image file into custom code endoSeg, see section below for code description, designed to extract cell features and fluorescence intensity characteristics at subcellular regions. Border masks were created using Procreate^®^ software, version 5.3.2, for iPad (Apple Inc.). Each mask was hand-drawn as a continuous brush stroke for all full-length cells in the field of view.

For analysis of visual identification of the calcium activity location in HAECs, the “time lapse differential” function in ZEN Blue 3.5 software was used to obtain only the area that depicts location of fluorescence intensity changes and, therefore, sites of Ca^2+^ spikes. The original files and resulting files from the time lapse differential were processed using “Maximal Projection” function in ImageJ, and the processed two images were merged.

To quantify the physical association of Caveolin-1 and Trpv4, the total number of the PLA puncta per cell were counted. Briefly, obtained images were processed using ImageJ (NIH) to subtract backgrounds and subsequent binary conversion. After binary conversion, PLA puncta were identified and were counted with the ‘analyze particles’ function.

Caveolin-1 cluster analysis was achieved by quantifying Caveolin-1 staining within individual cells as identified by VE-Cadherin boundaries. Each cell was further segmented into equal length thirds for the upstream, mid-body and downstream regions. Confocal images were processed using ImageJ (NIH) for binary conversion. Caveolin-1 clusters were quantified by measuring the number of particles and total area coverage using the ‘analyze particles’ function. The cluster index was then calculated using the following equation:

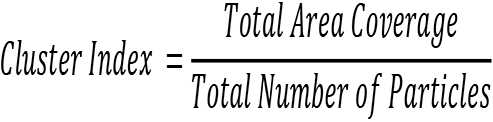

### endoSeg

To quantify the localization of fluorescence signal in cells, we employed a series of custom analysis routines, implemented in Python, termed endoSeg. First, a border mask is required to define the boundaries of all cells within the particular field of view (FOV). The mask is binarized using Otsu’s method and trimmed. Following segmentation, individual cells are passed through one of two separate but related analysis pipelines depending on whether the input data are multi-channel fluorescence images or time-series Ca^2+^ tracking images.

For multi-channel image sets, the individual cell masks from the watershed segmentation are used to calculate the orientation of the cell as well as additional parameters such as the centroid, area and aspect ratio. The centroid and orientation are then used to create a map that computationally splits the cell into three equally sized sectors, an upstream third, a middle third and a downstream third. The total fluorescence signal of the whole cell is then normalized to be zero mean and standard deviation one to remove the influence of cell-to-cell variation in absolute fluorescence intensity. The remaining negative values are set to zero. For each fluorescent channel of interest, the mean, maximum and Index of Dispersion (IoD) of the signal are calculated for each sector and recorded. Since the IoD is the ratio of the variance to the mean, the presence of a polarized signal is reported in the output based on the IoD value being over two in a specific region.

For time-series image sets, the analysis starts similarly, with orientation and sector maps being calculated. For each time-series, the mean fluorescence intensity is calculated from the whole cell and for each of the three sectors to give a one-dimensional plot of fluorescence intensity over time. The decay of fluorescence intensity due to bleaching is corrected using a Savitzy-Golay filter with a window size of 51 pixels and a 5th-order polynomial. These particular values were selected to produce a smoothed representation of the signal decay while ignoring the presence of intensity spikes to avoid over-subtraction. Following the intensity decay correction, a median filter is applied to remove any remaining spurious background and negative values are set to zero. The mean, maximum and IoD of the fluorescence time-series are calculated for the whole cell and each of the three sectors. We found that the IoD can be quite sensitive to the absolute fluorescence intensity of a particular cell. To make this value comparable across all cells within a given field of view, the IoD for each sector is divided by the IoD of the whole cell.

This ratio of IoD is then used to determine the presence of intensity spikes (representing rapid changes in fluorescent signal) with a value greater than 2 being used as the threshold. The sector in which the spikes are located is reported.

### Inflammatory Response

To test the role of Trpv4 activity on EC inflammatory response, HAECs were seeded into ibidi 0.4 μ-slides (ibidi #80176) and subject to flow-preconditioning using ibidi Pump System (ibidi #10902) for 48 h (unidirectional laminar flow, 20 dynes/cm^2^) then removed from flow. The flow-aligned HAECs were treated with the pro-inflammatory cytokine, TNF- α (10 ng/mL; R&D Systems #210-TA-020/CF) in the presence of DMSO or GSK1016790A (10nM; Sigma-Aldrich #530533) for 30 min. Inflammatory phenotype was characterized by quantifying NF-κB p65 nuclear localization and ICAM-1 expression as well as inflammatory gene expression signature by NFKBIA, VCAM1, CCL2, SELE, and TNF.

### Gene Expression Analysis

For qPCR analysis, total RNA was extracted from adherent cells using RNeasy Mini kit (QIAGEN #74104). RNA was converted to cDNA by reverse transcription and cDNA was quantified using a Nano-Drop 8000 (Thermo Fisher). Target genes were quantified using iTaq Universal SYBR Green Supermix (BioRad #1725125) and the appropriate primer pairs. Samples were run on a QuantStudio 6 Flex 384-well qPCR apparatus (Applied Biosystems). Each target gene was normalized to *HPRT*. Primer sequences (5′ – 3′) are listed below.

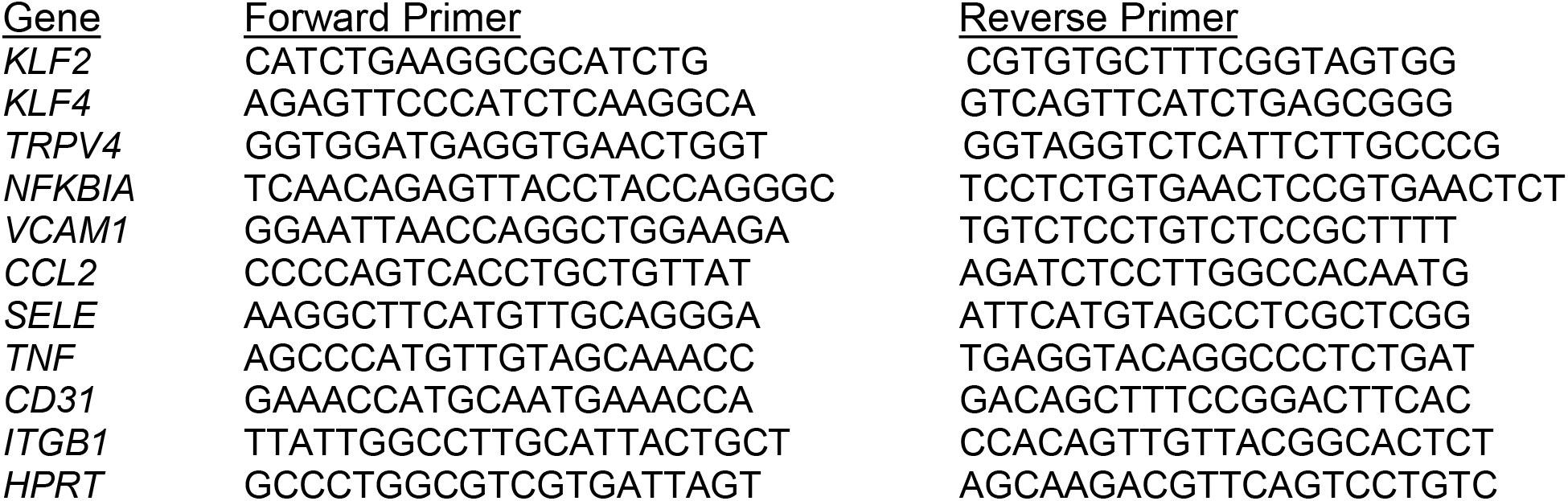

### Immunoblotting

To measure protein expression levels, HAEC monolayers were lysed in RIPA buffer (Fisher Scientific #PI89900) with Halt™ Protease and Phosphatase Inhibitor Cocktail (Thermo Scientific #78840) and 0.1 mM PMSF (Fisher Scientific #PI36978). Lysates were then centrifuged at 16,000g for 15 min at 4°C. Supernatants were collected and subjected to BCA protein assay (Pierce™ BCA Protein Assay Kit #23225) to quantify protein concentrations. Protein samples were run via sodium dodecyl sulphate–polyacrylamide gel electrophoresis (SDS-PAGE) and transferred to a polyvinylidene difluoride (PVDF) membrane using BioRad Trans-Blot® Turbo™ Transfer system (BioRad #1704150) and Trans-Blot® Turbo Mini PVDF transfer packs (BioRad #1704156). Subsequently, the membrane was incubated with 5% non-fat dry milk in Tris-buffered saline with Tween 20 (TBST) for 20 min at room temperature and then incubated overnight at 4°C with respective primary antibodies. The membrane was washed with TBST and incubated with HRP-conjugated secondary antibodies for 1 h. After washing the membrane was subjected to chemiluminescence method for visualization using ChemiDoc^TM^MP Imaging System (BioRad #17001402). Primary antibodies used for immunoblotting were as follows: α-Trpv4 (Alomone Labs #ACC-034), α-Total-eNOS (Santa Cruz #sc-376751), α- Caveolin-1 (Santa Cruz #sc-70516), α-GAPDH (Cell Signaling #2118S).

### Statistical Analysis

Statistical analysis was performed using GraphPad Prism software. The results are presented as mean ± SD. Depending on how many conditions were compared, either two-tailed independent *t*-test analysis or one-way analysis of variance (ANOVA) with Tukey’s post hoc multiple comparisons test was conducted. The Pearson correlation coefficient (r) was used to measure the strength of a linear association between two variables. P < 0.05 was considered statistically significant for all analyses.

### Drawings

Schematics in figures were created using BioRender (https://biorender.com).

## Code Availability

Custom code used for image analysis is available via public GitHub repository link: https://github.com/marcusgj13/endoSeg

## Acknowledgements

We thank M.T. Nelson (U Vermont) for helpful discussions, R. Wollman (UCLA) for providing GCaMP plasmids and J. Schwarz and M. Balles (ibidi GmbH) for providing the micropatterning slides. We thank the CNSI Nano & Pico Characterization Lab and M. Lake (UCLA) for assistance with AFM operation. The live cell 2-photon confocal imaging was performed in the UCLA Broad Stem Cell Research Center Imaging Core Facility. The Rosalind Franklin Institute is funded by UK Research and Innovation through the Engineering and 470 Physical Sciences Research Council. This work was funded by NIH grant P30 DK063491. S.H. was supported as a Jim Easton CDF Investigator; J.K. was supported by AHA Postdoctoral Fellowship 903306; P.T. was supported by P01 HL146358 and Leducq Foundation grant 19CVD04; J.J.M was supported by American Heart Association Career Development Award 19CDA34760007 and received a Justice Equity Diversity Inclusion award from the Life Science Editors Foundation, which provided editorial advice.

## Contributions

J.J.M. initiated, designed and supervised the study. S.H., J.W.A., J.K., E.C., M.W. and J.J.M. performed experiments. R.F., P.T. and P.T. shared reagents and experimental expertise. M.G-J. wrote analysis code and performed image analysis. P.T. and J.J.M wrote the paper.

## Corresponding author

Correspondence to Julia J. Mack.

## Supplemental Videos

For all videos, flow direction is left to right.

**Movie 1**. 30 min time-lapse recording of GCaMP expressing flow-aligned HAEC monolayer in the presence of high laminar flow (20 dynes/cm^2^).

**Movie 2**. 15 min time-lapse recording of GCaMP expressing flow-aligned HAECs in the presence of high laminar flow (20 dynes/cm^2^). Note, the cell outlined is used for analysis performed in Fig. 2b.

**Movie 3**. 15 min time-lapse recording of GCaMP expressing flow-aligned HAEC outlined for downstream tip (black) and cell body (white). Note that GCaMP fluorescence signal is displayed as Gold Lookup Table (LUT). Analysis of subcellular Ca^2+^ activity is presented in Fig. 2b.

**Movie 4**. 15 min time-lapse recording of GCaMP expressing HAECs in the presence of low flow (∼5 dynes/cm^2^).

**Movie 5**. 15 min time-lapse recording of GCaMP expressing HAECs in the presence of high flow (∼20 dynes/cm^2^).

**Movie 6**. 25 min time-lapse recording of GCaMP HAECs in flow convergence region of increasing shear stress from 5 dynes/cm^2^ to 20 dynes/cm^2^. Analysis of Ca^2+^ activity is presented in Fig. 3b.

**Movie 7**. 90 min time-lapse imaging of flow-aligned GCaMP HAEC exposed to 30 min laminar flow (20 dynes/cm^2^), followed by 30 min static (0 dynes/cm^2^) condition and then 30 min laminar flow (20 dynes/cm^2^). Note that GCaMP fluorescence signal is displayed as Gold Lookup Table (LUT). Analysis of Ca^2+^ activity at the downstream end is presented in Fig. 2f.

**Movie 8**. 10 min time-lapse imaging of GCaMP HAECs seeded on line-patterned chamber recorded in static conditions. Analysis of Ca^2+^ activity is presented in Supplemental Fig. 3d.

**Movie 9**. 30 min time-lapse recording of cholesterol depleted flow-aligned GCaMP HAECs in the presence of high flow (20 dynes/cm^2^). At 20 min, the Trpv4 agonist GSK10116790A (10 nM) is added to the culture flowing media. Ca^2+^ activity analysis is displayed in Fig. 4 d-f.

---

**Supplemental Fig. 1:**
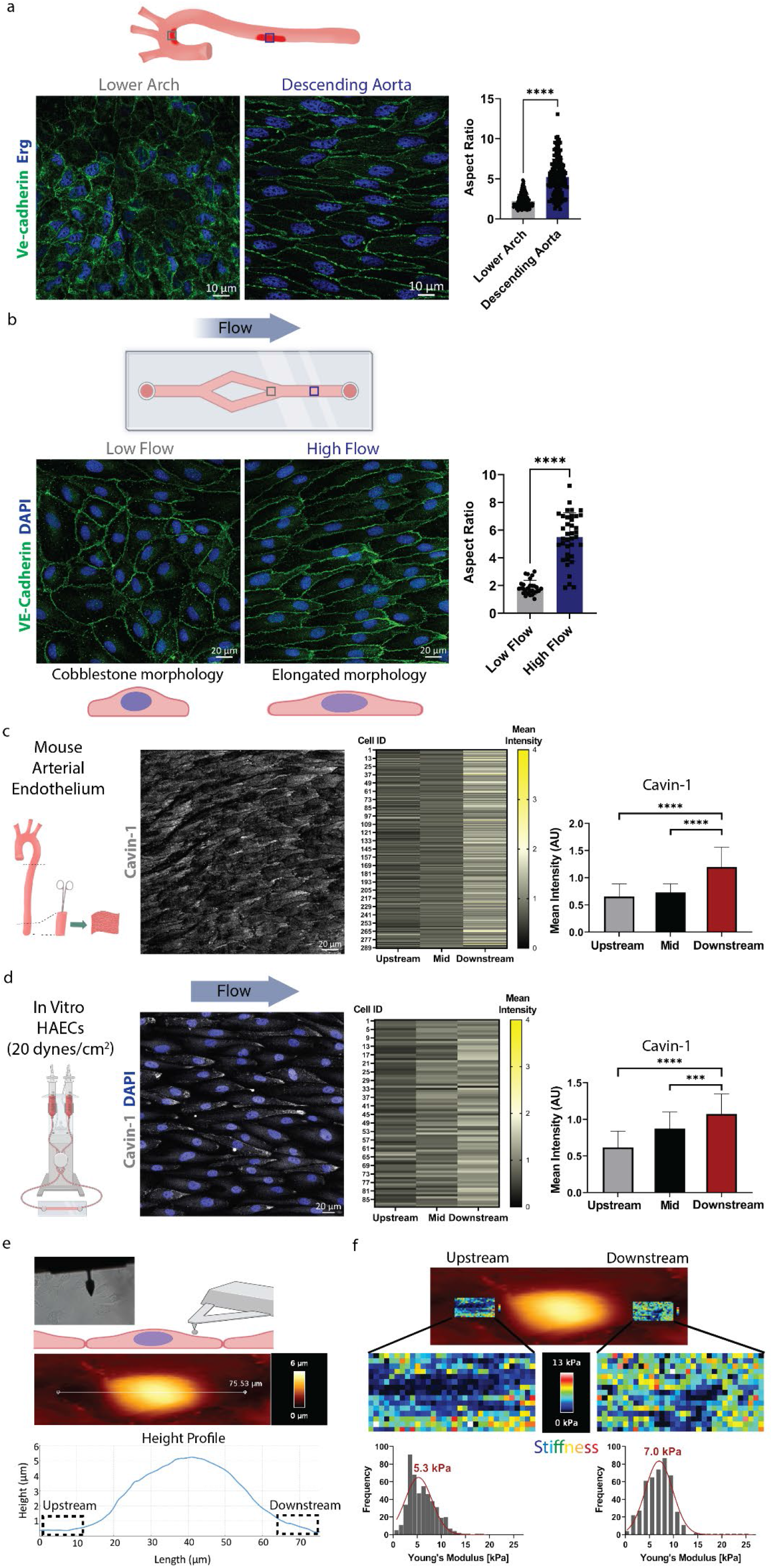
High flow induces arterial endothelial cell elongation and asymmetry of the cell surface. **a,** Representative images from *en face* imaging of the endothelium from a wildtype mouse aorta stained for Ve-cadherin and Erg1. Border masks were applied to calculate cell aspect ratio for 215 and 166 cells in the lower arch and in the descending aorta, respectively. \*\*\*\**P* < 0.0001 by two-tailed, unpaired *t* test. **b,** HAECs were seeded on y-shaped chambers and exposed to laminar flow for 48 h, then fixed and stained for DAPI and junctional marker ZO-1. Aspect ratio was quantified for cells in low-flow (∼5 dynes/cm^2^; n = 28) and high-flow (∼20 dynes/cm^2^; n = 38) regions. \*\*\*\**P* < 0.0001 by two-tailed, unpaired *t* test. **c,** Confocal imaging of Cavin-1 in the endothelium of wildtype mouse descending aorta. Individual cells were segmented into 3 equal-length regions (upstream, mid and downstream), and the staining intensity was determined for each segment in the number of cells indicated. Bar graph displays the mean ± SD with data analyzed by one-way ANOVA and post hoc Tukey’s multiple comparisons test. \*\*\*\**P* < 0.0001. **d,** As for (c) but using flow-aligned HAECs. \*\*\**P* < 0.001, \*\*\*\**P* < 0.0001 by two-tailed, unpaired *t* test. **e**, Atomic force microscopy (AFM) of flow-aligned HAEC surface: shown is the experimental setup with phase contrast image and the height profile of a representative cell. Upstream and downstream ends were clearly discernible. **f**, Cell stiffness was measured by calculating the Young’s modulus (kPa) from the force curves acquired at the upstream and downstream regions. The Young’s modulus values are displayed as a frequency plot with mean value for each region.

**Supplemental Fig. 2:**
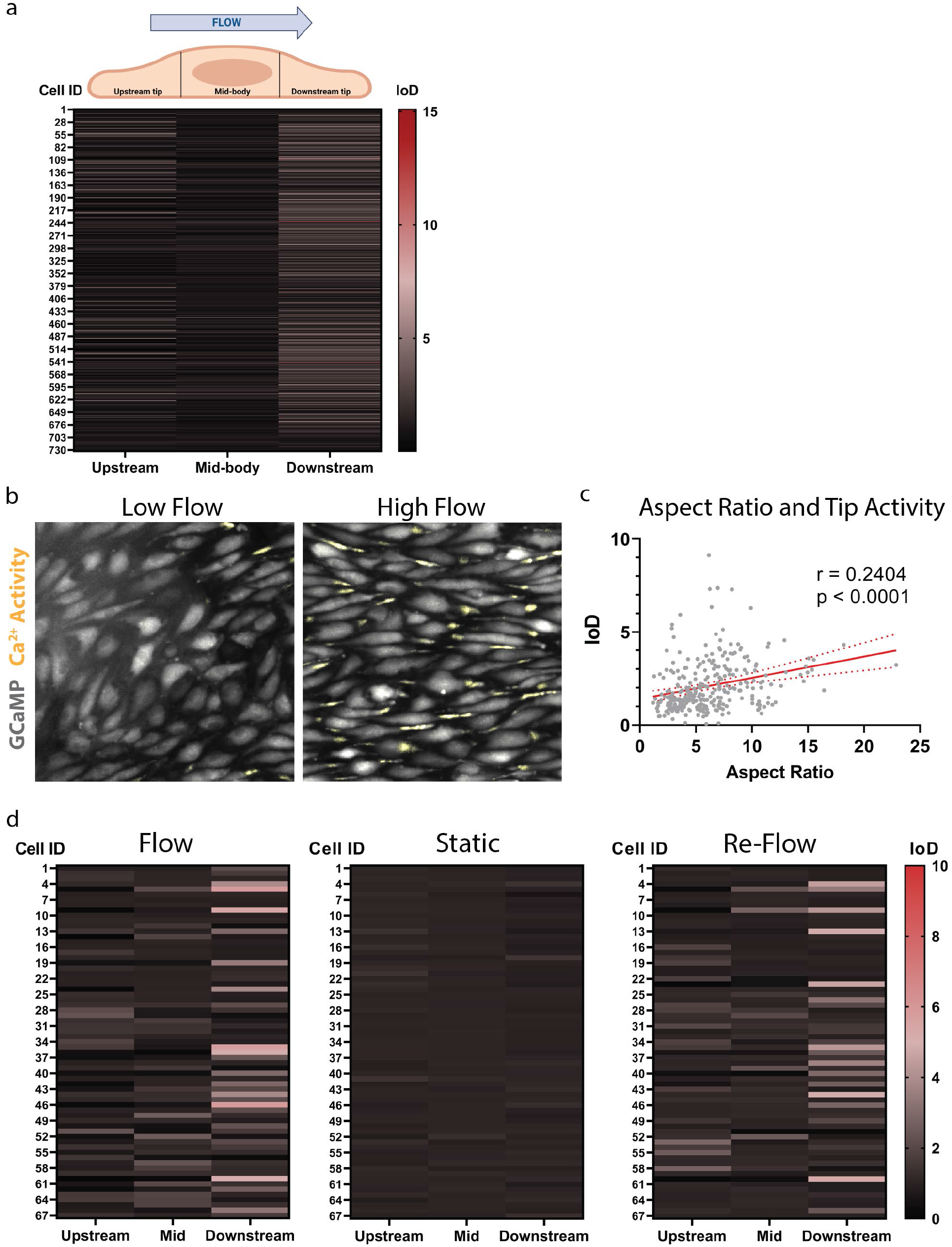
Ca^2+^ activity analysis in flow-aligned HAECs. **a,** Heat map displaying subcellular location of Ca^2+^ activity, plotted as the index of dispersion (IoD), for cells exposed to high laminar flow (20 dynes/cm^2^) and analyzed in Fig. 2c. **b,** Higher magnification images of cells in the low-flow and high-flow regions of the y-shaped slide in Fig. 2 where Ca^2+^ spikes are indicated in yellow. **c,** Correlation analysis of downstream end Ca^2+^ activity and cell aspect ratio. Pearson correlation coefficient (r) is used. **d,** IoD heat maps displaying subcellular location of Ca^2+^ activity for flow-aligned cells in the presence of high laminar flow (20 dynes/cm^2^) for 30 min, followed by no flow (0 dynes/cm^2^) for 30 min, and again exposed to high laminar flow (20 dynes/cm^2^) for 30 min. IoD plots show data from 67 cells across n = 3 biological replicates.

**Supplemental Fig. 3:**
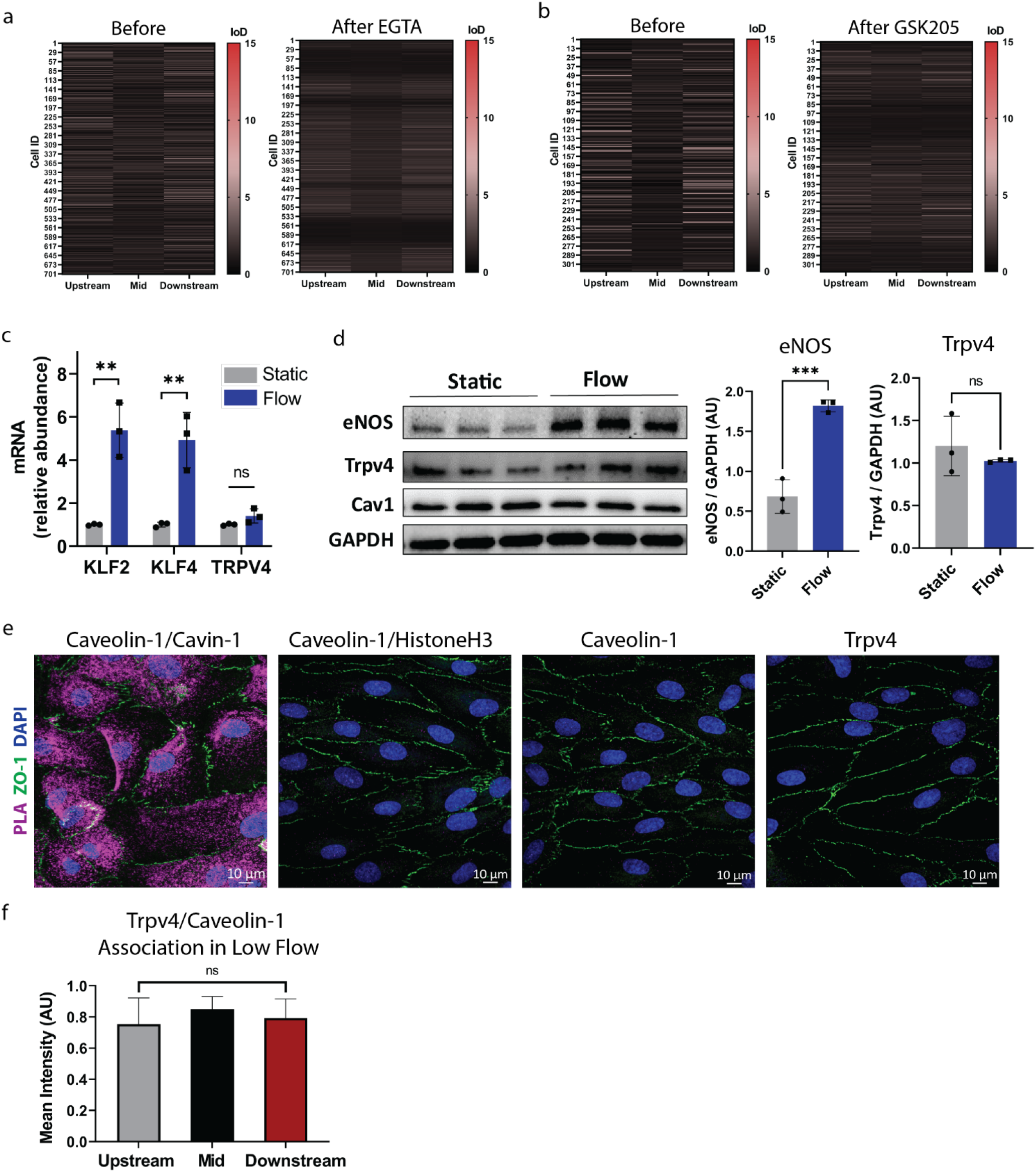
Trpv4 expression and localization under flow. **a,** Heat map displaying Ca^2+^ activity for cells in the presence of high laminar flow before and after treatment with EGTA (1.6 mM). **b,** Heat map showing the subcellular location of Ca^2+^ activity for cells under high laminar flow before and after treatment with GSK205 (20 µM). **c,** Gene expression analysis in HAECs cultured for 48 h in static or flow conditions shown as mean mRNA relative abundance ± SD (n = 3 biological replicates per condition). Note the flow-induced expression of KFL2 and KLF4 but unchanged expression for TRPV4. **d,** Western blot analysis of eNOS, Trpv4, Cav1 and GAPDH in HAECs cultured for 48 h under static or flow (20 dynes/cm^2^) conditions. Protein expression was determined by densitometry, normalized to GAPDH levels and then expressed as means ± SD (n = 3 biological replicates per condition). ns, not significant, \*\*\**P* < 0.001 by two-tailed, unpaired *t* test. **e,** Confocal images of representative controls for the PLA. Strong PLA was observed for Caveolin-1/Cavin-1. No PLA puncta were observed for Caveolin-1 and nuclear HistoneH3 or when Caveolin-1 or Trpv4 were used on their own. **f,** Quantification of the PLA for Trpv4/Caveolin-1 in HAECs in low-flow regions (as described for Fig. 3d). Shown are means ± SD (n = 49 cells analyzed) with statistics calculated using one-way ANOVA. ns, not significant.

**Supplemental Fig. 4:**
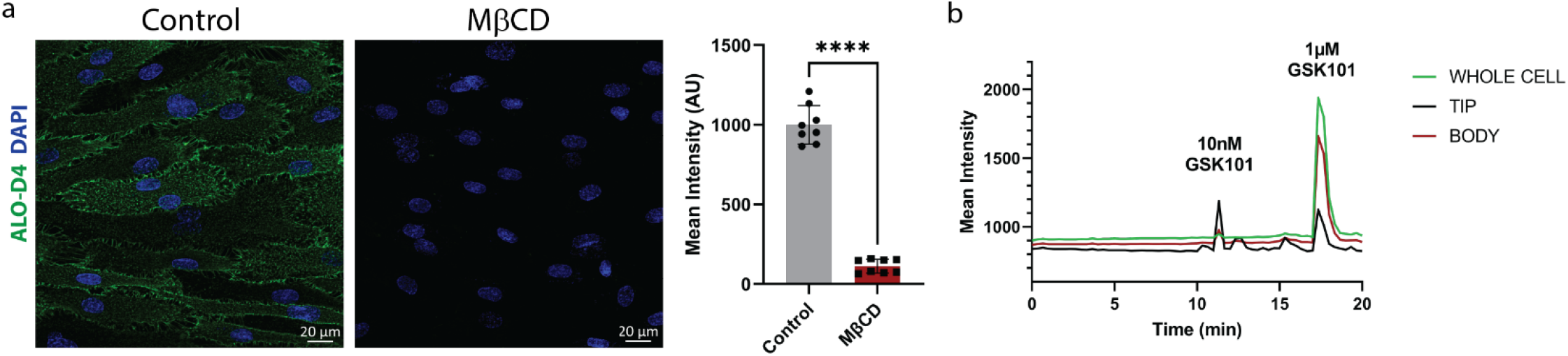
Cholesterol depletion using MβCD and induced Ca^2+^ activity. **a,** Flow-aligned HAECs were treated with or without MβCD (10 mM) in 1% LPDS statically for 30 min and then subjected to flow (20 dynes/cm^2^). After 2 h, cells were subjected to ALOD4-488 (20 µg/mL), fixed and then imaged. Shown are representative confocal images on the left and mean intensity ± SD of the ALOD4 fluorescent signal from 8 images per condition. \*\*\*\**P* < 0.0001 by two-tailed, unpaired *t* test. **b,** MβCD treated GCaMP HAECs were imaged under flow to record Ca^2+^ activity. No activity was recorded over 10 min. Addition of 10 nM GSK1016790A led to activity at the tip but not in the cell body. When 1 µM GSK1016790A was added, activity was observed throughout the entire cell.

**Supplemental Fig. 5:**
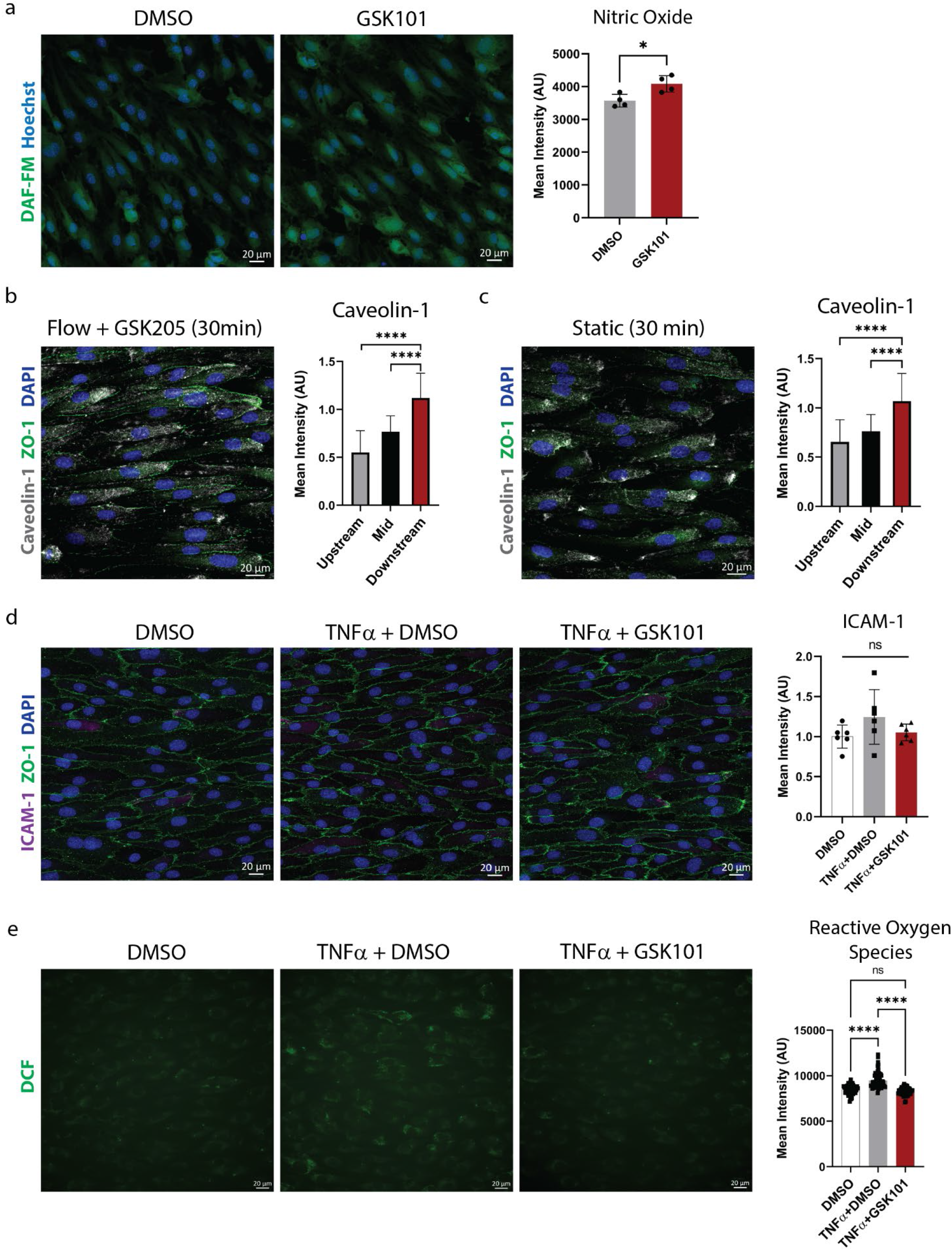
Trpv4 activation enhances NO and suppresses ROS production. **a,** Flow-aligned HAEC monolayers were treated with DMSO or GSK10116790A for 2 h followed by live cell imaging of DAF-FM and Hoechst. NO was quantified as the DAF-FM mean intensity signal ± SD (n = 4 biological replicates), and statistics were calculated using two-tailed, unpaired *t* test; \**P* = 0.0172. For (b-c), confluent monolayers of HAECs were exposed to laminar shear stress (20 dynes/cm^2^) for 48 h then treated for 30 min. After 30 min, monolayers were fixed and stained for Caveolin-1, ZO-1 and DAPI. Full-length cells were segmented into 3 equal-length regions, and Caveolin-1 staining intensity was quantified for each subcellular region. **b,** Representative image for flow-aligned HAECs treated with the Trpv4 antagonist GSK205 (20 µM) in the presence of laminar flow (20 dynes/cm^2^) for 30 min. Caveolin-1 staining intensity is quantified for 67 cells. Bar graph displays the mean ± SD with data analyzed by one-way ANOVA and post hoc Tukey’s multiple comparisons test. \*\*\*\**P* < 0.0001 (Upstream vs. Downstream; Mid vs. Downstream). **c,** Representative image for flow-aligned HAECs removed from flow for 30 min then fixed and stained. Caveolin-1 staining intensity is quantified for 61 cells. Bar graph displays the mean ± SD with data analyzed by one-way ANOVA and post hoc Tukey’s multiple comparisons test. \*\*\*\**P* < 0.0001 (Upstream vs. Downstream; Mid vs. Downstream). **d,** Flow-aligned HAEC monolayers were treated for 30 min in static conditions with DMSO or TNFα (10 ng/mL) in the presence or absence of GSK1016790A (10 nM). After 30 min, monolayers were fixed and stained for junctional marker ZO-1, inflammatory marker ICAM-1 and DAPI. ICAM-1 mean fluorescence intensity ± SD (n = 3 biological replicates with 2 images quantified per slide) did not show statistical difference across the groups. ns, not significant by two-tailed, unpaired *t* test. **e,** To measure reactive oxygen species (ROS), cells were loaded with CM-H_2_DCFDA (5 μM, DCF), then flow aligned for 48 h, followed by 1 h treatment in static conditions with DMSO, 10 ng/mL TNFα + DMSO or 10 ng/mL TNFα + 10 nM GSK1016790A. Mean DCF intensity signal ± SD is shown from 60 cells/condition. Data analyzed by one-way ANOVA and post hoc Tukey’s multiple comparisons test; \*\*\*\**P* < 0.0001 or not significant (ns).

